# Effects of innate immune activation by Toll-like receptor agonists on ethanol consumption and preference in FVB/NJ x C57BL/6J hybrid mice

**DOI:** 10.1101/2025.05.06.652465

**Authors:** Brent R. Kisby, Isabel Castro-Piedras, Sambantham Shanmugam, Igor Ponomarev

## Abstract

Excessive alcohol (ethanol) consumption is a hallmark of alcohol use disorder (AUD). Activation of innate immune system and proinflammatory signaling in the brain may play a key role in promoting alcohol consumption and development of AUD in humans. Innate immune activation by toll-like receptor (TLR) agonists in rodents is associated with release of proinflammatory cytokines and changes in alcohol consumption, and these effects are genotype– and sex-dependent. For example, C57BL/6J male, but not female mice increase alcohol intake after TLR3 activation. In order to better understand the interactions between neuroimmune signaling, genotype, and sex and their effects on ethanol drinking, males and females of more genotypes need to be tested. The goal of this study was to test the effects of innate immune activation on ethanol consumption and neuroimmune molecular profiles of F1 hybrid mice from reciprocal crosses between C57BL/6J (B6) and FVB/NJ (FVB) mouse strains, which are animals with high levels of ethanol intake. Animals were randomly assigned to receive intraperitoneal injections of either saline, Poly(I:C) (PIC, 2 or 10 mg/kg), a TLR3 agonist, or lipopolysaccharide, (LPS, 0.1 mg/kg), a TLR4 agonist, administered every 4 days for a total of 10 injections and subjected to a 2-bottle choice every-other-day ethanol drinking paradigm for a total of 18 dinking sessions, which generated high levels of voluntary ethanol consumption. Six and 24 hours after the last injection, brains were removed, frontal cortex dissected, and levels of 3 proinflammatory cytokines (*Tnfa*, *Il1b*, *Ccl5), as well as Tlr3*, and *Tlr4* were measured using qPCR. Immune activation by PIC produced escalation of ethanol drinking, while LPS resulted in a reduction of ethanol consumption or a trend to reduce drinking in males but not females of both FVB/B6 and B6/FVB crosses. Furthermore, activation of TLR3 by PIC produced sex-specific time course responses of pro-inflammatory cytokines, which may, at least in part, explain behavioral differences. Taken together, these results validate previous findings that the effects of immune activation on ethanol consumption depend on genotype, sex, and mode of activation (TLR3 vs TLR4) and suggest that FVB/B6J and B6J/FVB F1 males are a suitable model to study TLR3-dependent escalation of alcohol drinking.

**Highlights:**

- Immune activation by Toll-like receptor 3 (TLR3) agonist, Poly(I:C), produced an escalation of ethanol drinking, while immune activation by TLR4 agonist, LPS, reduced ethanol intake in male but not female FVB/NJ x C57BL/6J hybrid mice.
- Poly(I:C)-induced escalation of alcohol consumption in males was reproducible and consistent across different Poly(I:C) doses.
- Activation of TLR3 by Poly(I:C) produced sex-specific time course responses of pro-inflammatory cytokines, which may, at least in part, explain sex differences in alcohol consumption.
- Our data suggest that FVB/NJ x C57BL/6J hybrid male mice are a suitable model to study TLR3-dependent escalation of alcohol drinking.

## Introduction

Neuroimmune signaling refers to immune-related processes within the central nervous system (CNS) and their interactions with the peripheral immune system. Alterations in neuroimmune signaling have been linked to a broad spectrum of brain disorders including alcohol use disorder (AUD) (Crews et al., 2015; Cui et al., 2014; Mayfield & Harris, 2017; Perkins et al., 2019). Activation of the innate immune system results in the release of pro-inflammatory cytokines, which may lead to behavioral abnormalities including high alcohol intake, an AUD hallmark (Blednov et al., 2011; Hitzemann et al., 2021; McCarthy et al., 2018; Mulligan et al., 2006). According to the neuroimmune hypothesis of “alcohol addiction”, drinking alcohol promotes pro-inflammatory signaling, which, in turn, results in more alcohol intake, creating a positive feedback loop that may lead to excessive alcohol consumption (Mayfield & Harris, 2017). There is a lack of understanding of how the innate immune system modulates the transition from low to high drinking, how changes in immune signaling are integrated into neural networks, and how disease progression is orchestrated by multiple neuroimmune components. Another critical gap in knowledge is the role of sex and genetic factors in these processes.

Rodent models have been widely used to study the role of neuroimmune signaling in the transition from low or moderate alcohol consumption to excessive drinking. Recent studies showed that activation of the innate immune system by administration of small molecule ligands of Toll-like receptors (TLRs) affects alcohol consumption in a sex-, genotype-, and TLR type-dependent manner (Blednov et al., 2011; Giffin et al., 2022; Hitzemann et al., 2021; Lovelock et al., 2022; Warden et al., 2019a, 2019b; Warden, DaCosta, et al., 2020; Warden, Triplett, et al., 2020). For example, administration of synthetic double-stranded RNA, Poly(I:C) (PIC), a TLR3 agonist, resulted in escalated alcohol consumption in C57BL/6J male but not female mice (Warden et al., 2019a, 2019b), while bacterial lipopolysaccharide (LPS), a TLR4 agonist, caused genotype-specific increases in alcohol intake in different mouse strains (Blednov et al., 2011). Studies suggest that these behavioral differences may, at least, in part, be explained by different immune responses followed by innate immune activation (Hitzemann et al., 2021; Silva-Gotay et al., 2021; Warden, DaCosta, et al., 2020; Warden et al., 2019a, 2019b). For example, C57BL/6J females had a delayed and longer-lasting elevation of pro-inflammatory cytokines after TLR3-dependent immune activation than male mice, which may explain sex differences in PIC-induced changes in alcohol drinking (Warden et al., 2019a, 2019b).

To further understand the role of innate immune activation and its interactions with sex and genotype in regulation of alcohol consumption we tested male and female F1 hybrid mice from reciprocal crosses between C57BL/6J (B6) and FVB/NJ (FVB) mouse strains. These hybrid mice consume more ethanol (EtOH) than B6 or FVB strains alone (Agrawal et al., 2014; Blednov et al., 2005, 2010; Ozburn et al., 2010) and may serve as a suitable model to study the effects of innate immune activation on alcohol drinking. We tested the effects of different doses of TLR3 and TLR4 agonists on chronic EtOH consumption and neuroimmune molecular profiles and compared findings with published data. Our results presented below further suggest that the effects of immune activation on EtOH consumption depend on genotype, sex, and mode of activation and could, at least, in part, be explained by temporal profiles of neuroimmune molecules.

## Materials and methods

### Animals

EtOH-naïve female and male C57BL/6J (B6) (cat: 000664; Jackson Labs) and FVB/NJ (FVB) (cat: 001800; Jackson Labs) mice were ordered at 7 weeks of age and allowed 1 week of habituation in the breeding colony facility prior to breeding. Generation of the F1 crosses have been previously reported (Blednov et al., 2005, 2010; Ozburn et al., 2010). Briefly, animals were paired with a single female to a single male. To generate FVB/B6 F1 mice, FVB females were bred with B6 male and for B6/FVB F1 mice, B6 females were bred with FVB males. Experimental animals were weaned at 21 days of age and were transferred to the experimental room for at least 1 week prior to behavioral testing. Animals used in this study were aged between 10-12 weeks of age at the start of behavioral experiments. Animals for experiment 1 were split across 2 cohorts (cohort 1, n=48; cohort 2, n=68). We did not observe statistically significant differences between the cohorts (results not shown) and data for the two cohorts were combined. Animals in experiment 2 were tested in a single cohort (n=31). Prior to the initial injection, animals were counter-balanced for age, litter, treatment, and sex for the respective treatment groups and shelves within the experimental room. Animals were on a normal 12:12 hour light cycle with lights on at 6 am. Animals were given *ad libitum* food (PicoLab Rodent Diet 20; St. Louis MO) and water. Injections, bottle change, and weighing were performed between 9:30 am and 11:00 am. All procedures were approved by the Texas Tech University Health Sciences Institutional Animal Care and Use Committee (IACUC) (Protocol # 19009).

### Alcohol Consumption

We utilized the 2-bottle choice (2-BC) Every-other-day (EOD) drinking paradigm which has been previously shown to progressively escalate EtOH consumption in mice (Blednov et al., 2005, 2010; Ozburn et al., 2010; Warden et al., 2019a, 2019b). Briefly, animals were presented with either a single 50 ml conical bottle of water or a single bottle of 15% v/v diluted EtOH solution. EtOH was diluted from 190 proof (Pharmco, Shelbyville, KY) into distilled water. Each conical sipper tube used a #6 rubber stopper duel-ball bearing sipper. For a total of 24 hours, animals were allowed voluntary access to either water or EtOH. After 24 hours, the bottles were removed and replaced with 2 bottles of distilled water for 24 hours. EtOH bottle was initially presented on the right side and the EtOH bottle was switched between EtOH sessions to prevent side preference. EtOH consumption was measured as g/kg/24 hours. Two sessions of 24 hours were averaged for statistical analysis.

### Reagents

#### LPS

We used the O111:B4 strain of LPS (Sigma Aldrich, Burlington, MA, Lot# 0000137286, Cat# L3024-25mg), which was used in other EtOH drinking studies (Blednov et al., 2011; Decker Ramirez et al., 2023) and showed robust immune response. We prepared a stock solution with concentration of 1 mg/ml using endotoxin-free 0.9% saline (Quality Biological; Gaitensburg, MD). Prior to behavioral testing, we confirmed that LPS induced immune response in the frontal cortex in a separate set of animals by measuring cytokine levels at 6 hours after i.p. injection (data not shown). Aliquots of 100 ul were stored in –20° C until injection day. The remaining drug was discarded.

#### PIC

High Molecular Weight (HMW) Poly(I:C) (PIC) (Cat #: *Tlr3*-pic-5; Invivogen, San Diego, CA) was prepared using standard manufacturing protocol from Invivogen. Briefly, PIC was prepared by resuspending the lyophilized pellet with pyrogen-free saline. Solution was heated in a bead bath at 67 degrees for 10 minutes and let rest for 1 hour prior to aliquoting. Aliquots were stored at –20 degrees until the day of injection. Any remaining drug was discarded.

### RNA Isolation, cDNA preparation, and qPCR

Total RNA was isolated from mouse frontal cortex at the end of experiment 1 using the TRIzol reagent according to the manufacturer’s recommendations (Invitrogen, Carlsbad, CA, USA). 2.5 µg total RNA was reverse-transcribed using the Applied BioSystems cDNA synthesis kit in a 20 µL reaction and diluted at a 1:2 ratio with Nuclease free water. Real time PCR amplification was performed on a Biorad CFX384 Real time system (Hercules, CA). A 10 µL real time qPCR reaction containing 2 µL of the cDNA, 5 µL of TaqMan Universal Master Mix (Applied Biosystems, Bedford, MA, USA), 20 pmol of the respective primer/probe mix was used to determine the mRNA expression (Table: 1). PCR cycling conditions included an initial denaturation step at 95°C for 30s followed by 40 PCR cycles at 95°C for 5s and 60°C for 30s. The expression of the target gene was determined relative to the geometric mean of the following housekeeping genes: *Rn18s* and *Malat1*. Relative amplification of each transcript was further calculated as fold difference from untreated samples using the comparative 2^ΔΔ^-Ct method (Schmittgen & Livak, 2008).

### TaqMan probes

| Gene symbol | Gene name | Accession number | Dye |
| --- | --- | --- | --- |
| *Tnf* | tumor necrosis factor | Mm00443258 | FAM |
| *Il1b* | interleukin 1 beta | Mm00434228 | FAM |
| *Tlr3* | toll-like receptor 3 | Mm01207404 | FAM |
| *Tlr4* | toll-like receptor 4 | Mm00445273 | FAM |
| *Ccl5* | chemokine (C-C motif) ligand 5 | Mm01302427 | FAM |
| *Rn18s* | 18S ribosomal RNA | Mm03928990 | FAM |
| *Malat1* | Metastasis-associated lung adenocarcinoma transcript 1 | Mm01227912 | FAM |

### Experimental Design

*Experiment 1*. *Effects of PIC and LPS on EtOH consumption and preference as well as expression of pro-inflammatory genes in FVB/B6 and B6/FVB mice*. Animals were randomly assigned to receive intraperitoneal injections of either saline, PIC (10 mg/kg) or LPS, (0.1 mg/kg), administered every 4 days for a total of 10 injections. Saline control animals were given a per-body weight equivalent volume. Twenty four hours after the first injection, animals were introduced to their first 2-BC EOD exposure (see details above) for a total of 18 sessions. Six or 24 hours after the last injection, animals were sacrificed, brains harvested and frontal cortex (estimated between Bregma +1.09 to +2.9) dissected for the analysis of immune signaling gene expression. Tissues were snap frozen in liquid nitrogen and stored at –80°C until further RNA extraction.

*Experiment 2: Effects of different doses of PIC on EtOH consumption and preference in FVB/B6 mice*. Similar to experiment 1, animals were counterbalanced based on litter and assigned to receive intraperitoneal injections of either saline or PIC (2 or 10 mg/kg) administered every 4 days for a total of 10 injections. The 2-BC EOD EtOH drinking paradigm was used as described above.

### Statistics

For behavioral data from experiment 1 we used a 2-way mixed design analysis of variance (ANOVA) using Prism (Graphpad version 10; San Diego, CA). The behavioral drinking data were presented and analyzed as an average of two drinking sessions within an injection cycle. Our analysis consisted of drug Treatment (saline, LPS, and PIC) being the between-subject factor and EtOH Sessions being the within-subject factor. We used the Tukey post-hoc test to correct for multiple comparisons. Because of baseline sex differences in EtOH consumptions, males and females were analyzed separately. In experiment 2, we used a similar 2-way mixed design ANOVA. A statistically significant Treatment x EtOH Sessions interaction was followed by a one-way ANOVA for the last 2 EtOH sessions, followed by a one-tailed Dunnett post-hoc test to compare different doses of PIC to control.

To analyze qPCR data in experiment 1, we used 2-way ANOVA with Tukey post-hoc test. For figure 3 data, the between-subject factors were Timepoint (6 and 24 hours) and Genotype (FVB/B6 and B6/FVB). Males and females were analyzed separately. Statistical analysis used the ΔΔCt values and data were shown as 2^-ΔΔ^Ct values. Because we did not observe a robust effect of genotype, we combined the genotypes and performed a 2-way ANOVA with Timepoint and Sex as between-subject factors. This analysis was performed for the effects of PIC only, because of a more robust gene expression response, compared to the effects of LPS (fig. 4). A statistically significant effect was determined to have a p-value of less than 0.05. Data is reported as mean ± standard error of the mean (SEM).

**Figure 1:**
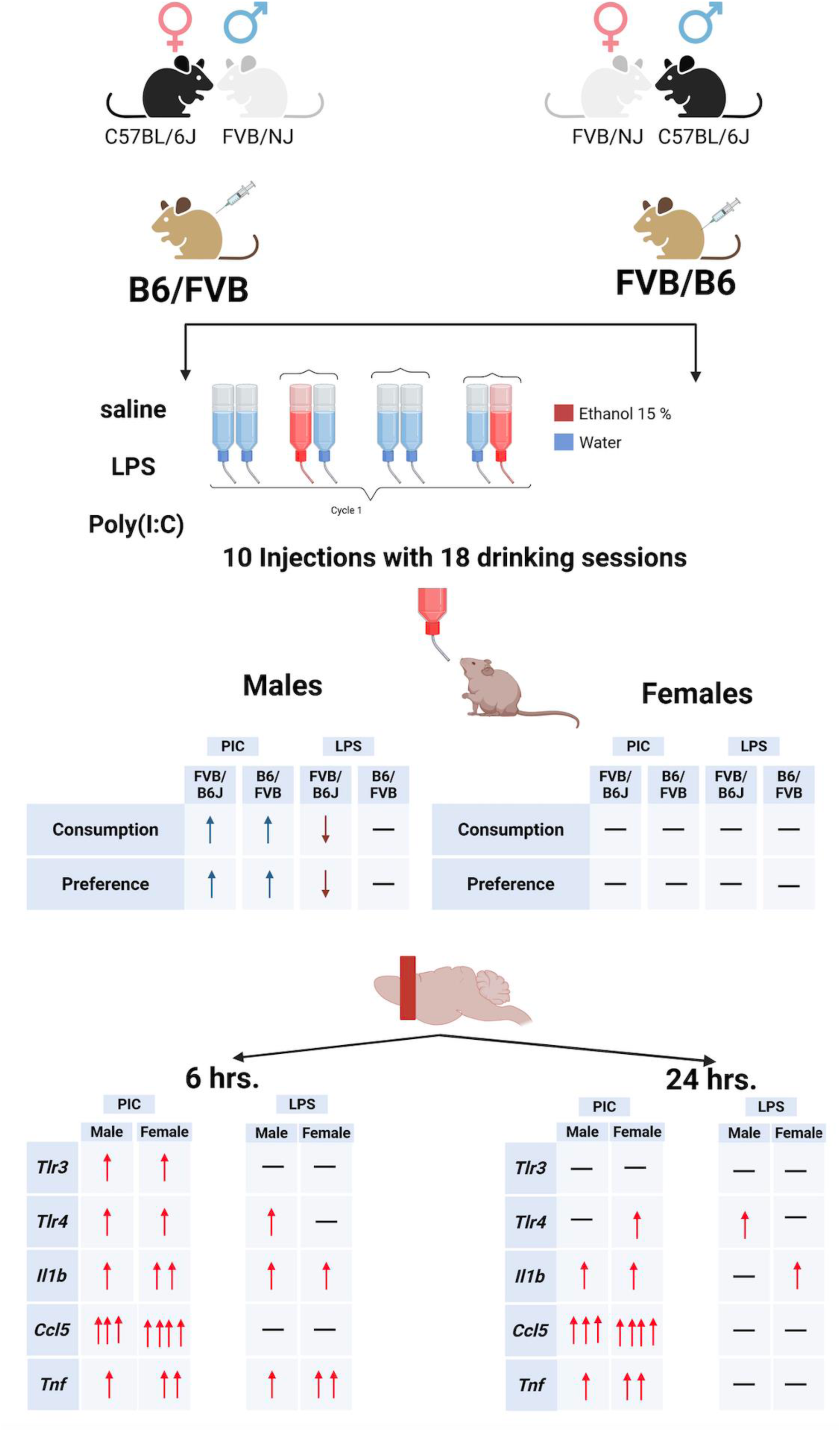
Graphical abstract showing experimental design and main results.

**Figure 2:**
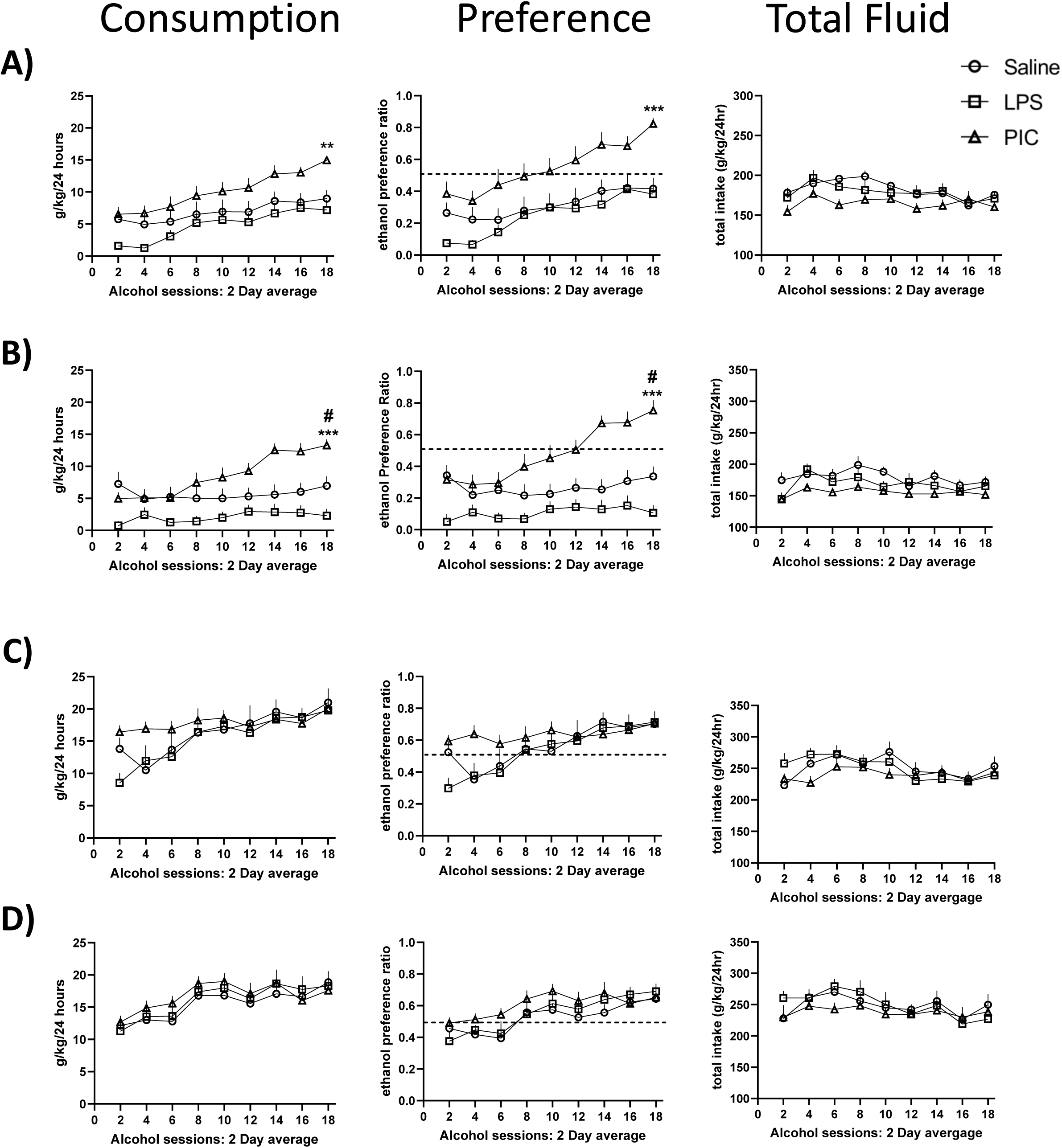
Behavioral results from Experiment 1. EtOH consumption (left panels), preference (middle panels) and total fluid intake (right panels) in male B6J/FVB (A), male FVB/B6J (B), female B6J/FVB (C), and female FVB/B6J (D) hybrid mice. Saline versus PIC at the final 2-day average alcohol sessions (*: p<0.05; **: p<0.01; ***: p<0.001); Saline versus LPS at the final 2-day average alcohol sessions (#: p<0.05). Error bars are SEM.

**Figure 3.**
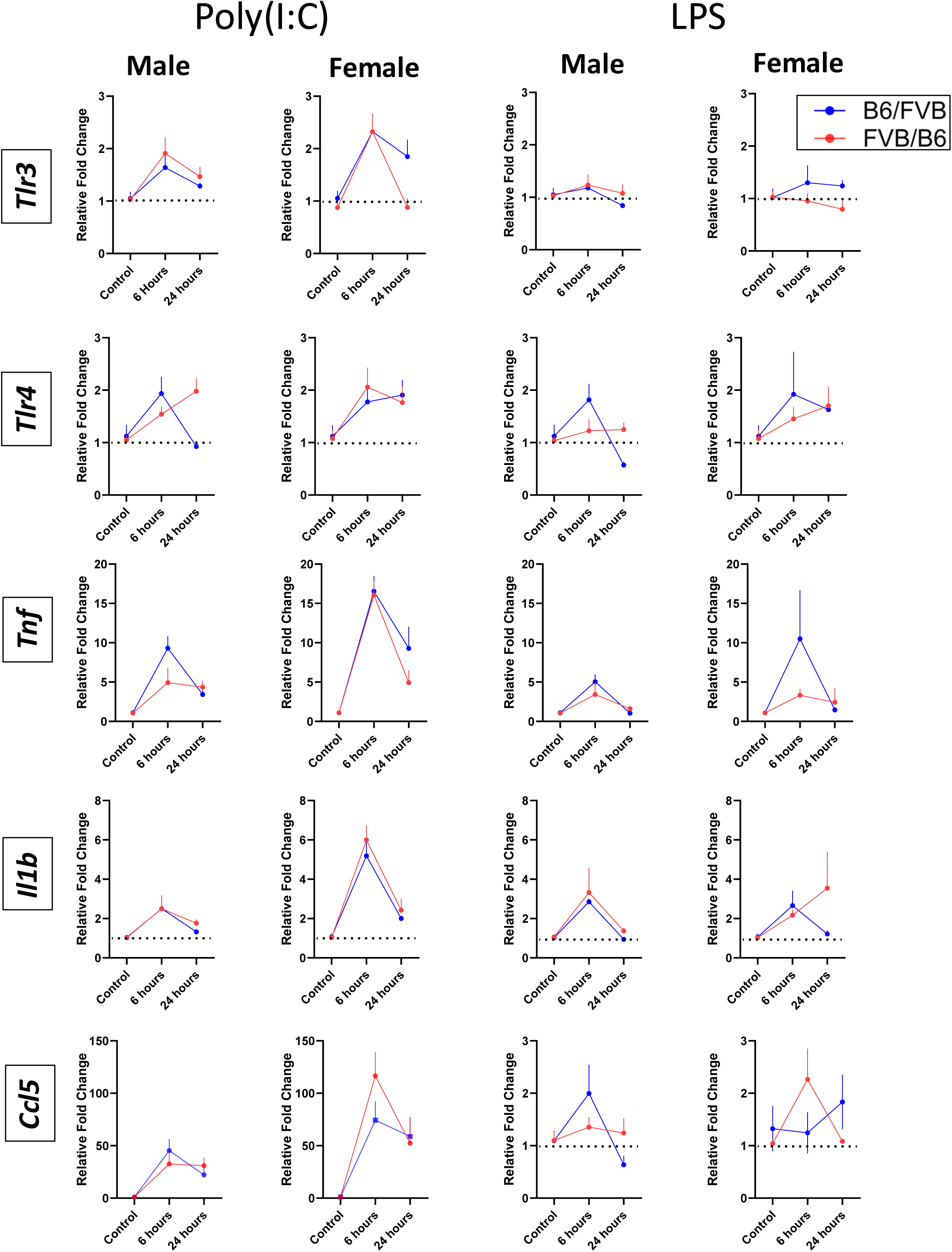
Gene expression results from Experiment 1. The mRNA levels of the indicated genes were measured by qPCR. Levels of each transcript were normalized to geometric mean of 18s and Malat1 and saline treated groups (fold-change ± SEM). Data is represented as 2^-ΔΔCt^ and statistics were performed on the ΔΔCt values. Error bars are SEM.

**Figure 4.**
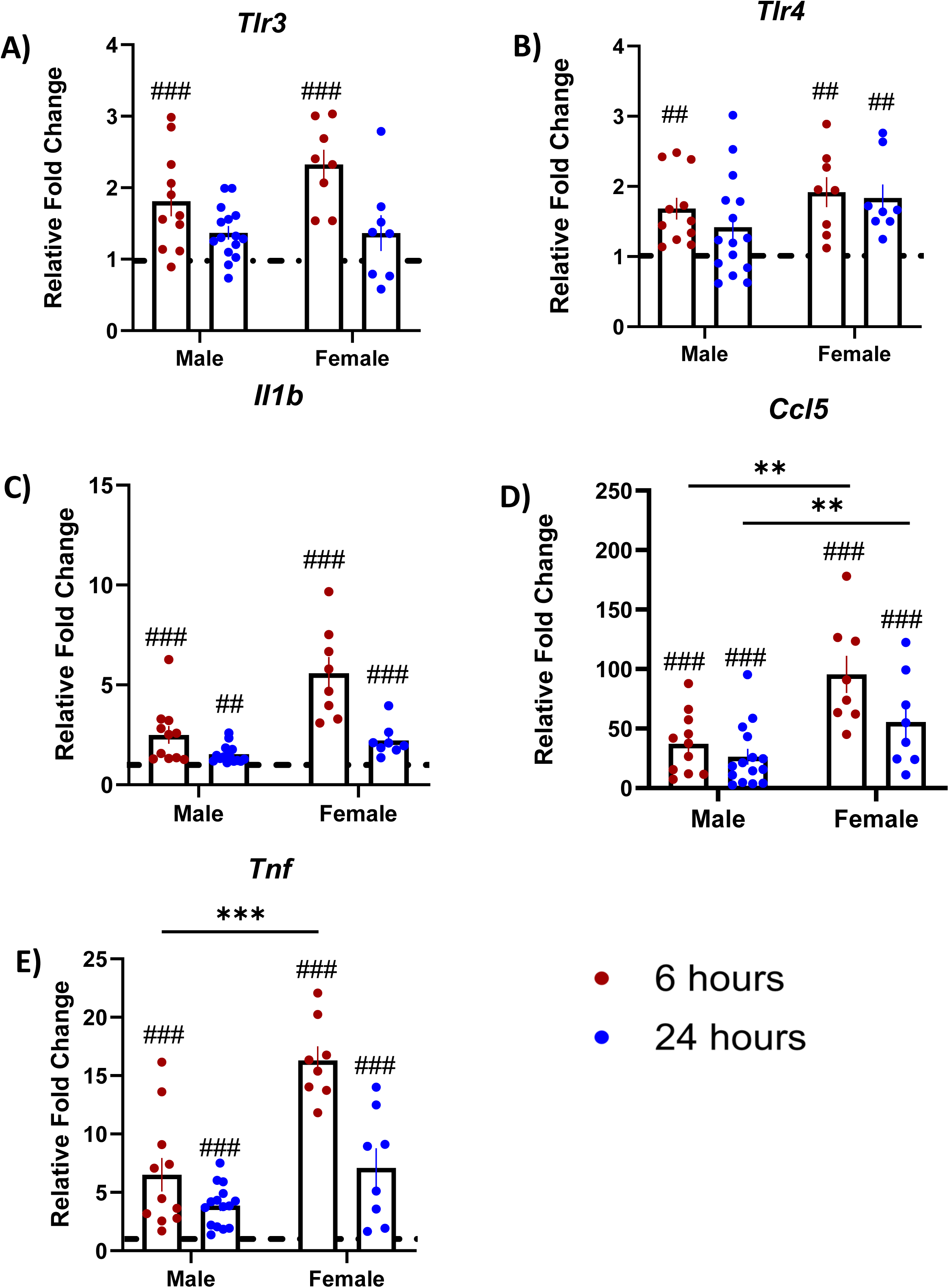
Gene expression results after PIC from Experiment 1 with combined data across genotypes. Shown are PIC-induced fold changes relative to saline. Two-way ANOVA was followed by post-hoc tests at each time point. PIC-induced changes (#: p<0.05; ##: p<0.01; ###: p<0.001; ####: p<0.0001). Sex differences (*: p<0.05; **: p<0.01; ***: p<0.001). Error bars are SEM.

## Results

### Experiment 1: Behavioral data

Behavioral data are shown in figure 2. Details of statistical analysis are shown in supplemental table 1. Treatment with either the TLR3 agonist, PIC, or the TLR4 agonist, LPS, resulted in changes in alcohol consumption in male but not female mice, as we observed a main effect of Treatment in B6/FVB (fig. 2A, left panel) and FVB/B6 males (fig. 2B, left panel), but not B6/FVB (fig. 2C, left panel) or FVB/B6 (fig. 2d, left panel) females. Post-hoc comparisons showed that male mice of both crosses treated with PIC consumed more EtOH than control mice, while LPS-treated FVB/B6 males consumed significantly less EtOH than saline-treated mice. There was a main effect of EtOH drinking Sessions in both crosses of both sexes, indicating that mice generally consumed more EtOH over time. We also observed a statistically significant interaction of Treatment x EtOH Session in FVB/B6 male mice. This effect was mainly driven by a sharper increase in consumption in PIC-treated mice, compared to control animals. Post-hoc comparisons showed that PIC-treated mice consumed significantly more ethanol than saline-treated animals during the last 2 drinking sessions. Analysis of EtOH preference data revealed the same group effects as were found for EtOH consumption (fig. 2A-D; middle panels). Treatment with either drug did not change total fluid intake across sex and genotype (fig. 2A-D; right panels).

### Experiment 1: Gene expression data

Molecular data are shown in figures 3 and 4. For details of all statistical results, see supplemental table 1. Two-way ANOVAs with Timepoint and Genotype as between-subject factors revealed no main effects of Genotype for either gene, except for *Tlr3* for females after either PIC or LPS (fig. 3). On the other hand, majority of main effects of Timepoint were statistically significant after either PIC or LPS, with the 6-hour timepoint showing consistently higher expression after immune activation by PIC compared to control. Twenty four hours after PIC injection, *Tnf* and *Ccl5* were still elevated in both sexes and genotypes. Statistically significant Genotype by Timepoint interactions were only detected for *Tlr4* and *Tnf* in males.

Two-way ANOVAs with Timepoint and Sex as between-subject factors reveled main effect of Timepoint for all genes as well as main effect of Sex and statistically significant Timepoint by Sex interactions for *Il1b, Tnf* and *Ccl5* (fig. 4). Both males and females showed a robust increase in gene expression 6 hours after PIC injection and a relative decrease at the 24-hour timepoint. Timepoint by Sex interactions for the three pro-inflammatory molecules were mainly driven by a sharper increase in expression at 6 hours and a slower decrease at 24 hours in females compared to males.

### Experiment 2: Behavioral data

The goal of this experiment was replication of results observed in experiment 1 for PIC and, also testing a lower dose of PIC (fig. 5). For details of statistical results, see supplemental table 1. For EtOH consumption, there was no main effect of Treatment in either males or females (fig. 5A-B), left panels). There was a statistically significant effect of EtOH Sessions in both sexes and a statistically significant Treatment x EtOH Sessions interaction in males only. This interaction was followed by a one-way ANOVA on the final two-day averaged drinking sessions, which was statistically significant. One-tailed Dunnett post-hoc tests revealed that both PIC doses resulted in an increase in EtOH consumption, compared to control. For EtOH preference, there were main effects of Treatment, EtOH Sessions, and a Treatment x EtOH Sessions interaction in males (fig. 5A, middle panel) and only a main effect of EtOH Sessions in females (fig. 5B, middle panel). One-way ANOVA on the final two-day averaged drinking sessions followed by one-tailed Dunnett post-hoc tests revealed that both PIC doses resulted in an increase in EtOH preference, compared to control. For total intake, there was a main effect of Treatment in both sexes, driven by a decrease in total intake after the higher dose of PIC (fig. 5A-B, right panels).

**Figure 5.**
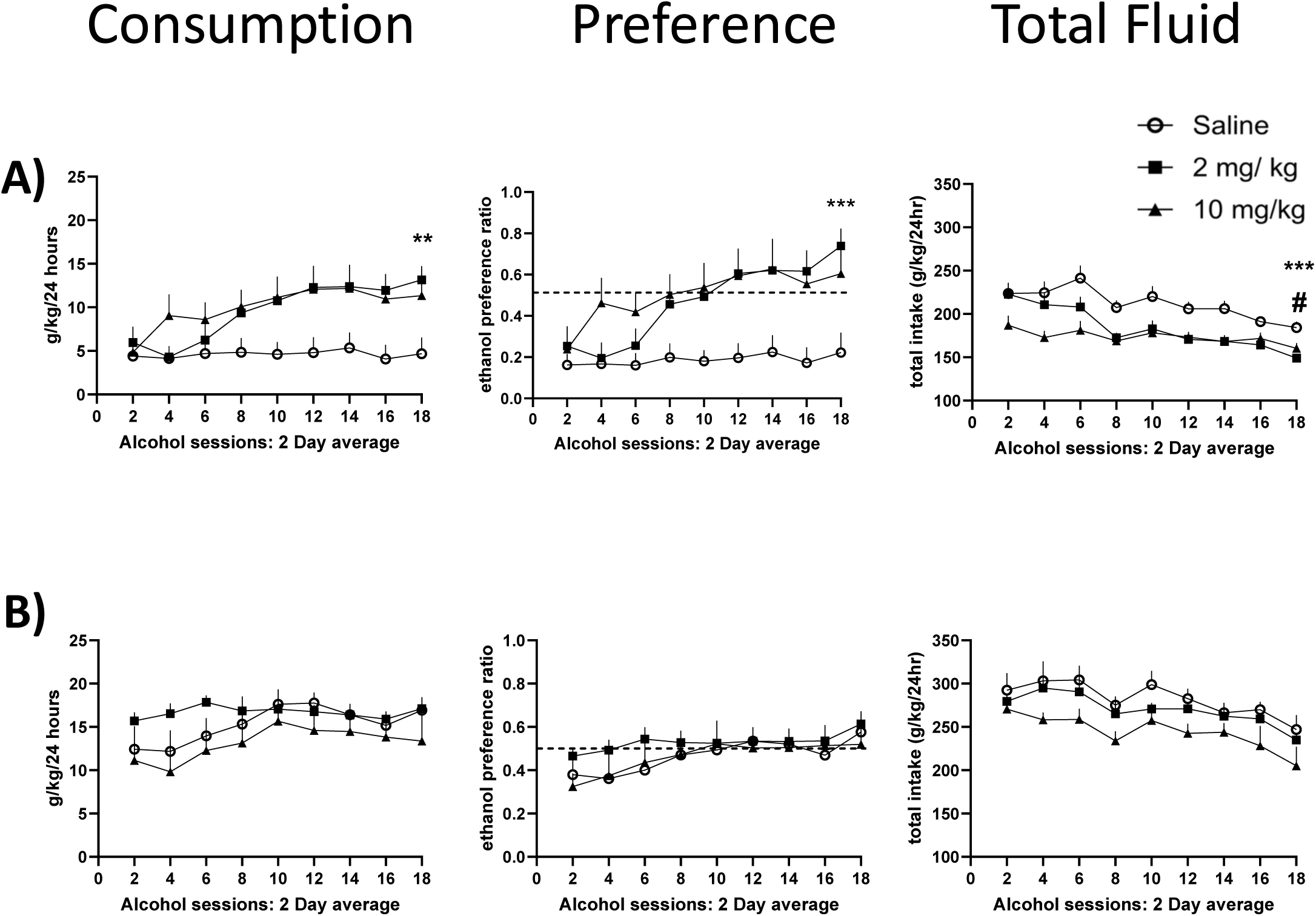
Behavioral results from Experiment 2. EtOH consumption (left panels), preference (middle panels) and total fluid intake (right panels) in male FVB/B6J (A) and female FVB/B6J (B) hybrid mice. Saline vs. 2 mg/kg at the final 2-day average alcohol sessions (**: p<0.01; ***: p<0.001). Control vs. 10 mg/kg at the final 2-day average alcohol sessions (#: p<0.05). Error bars are SEM.

## Discussion

The main goal of this study was to test the effects of innate immune activation on EtOH consumption and preference in male and female F1 hybrid mice from reciprocal crosses between C57BL/6J and FVB/NJ mouse strains. We tested the general hypothesis that TLR-dependent immune activation in male and female F1 hybrid mice would result in changes in EtOH consumption and preference. We showed that females did not change their ethanol consumption or preference in response to immune activation, while males were more responsive, with a TLR3-dependent increase in EtOH drinking in both hybrid genotypes and a TLR4-dependent decrease in EtOH intake in FVB/B6 mice. Our experiment 2 showed that the TLR3-depepndent effects were robust and could be reproduced using the same or a smaller dose of PIC. A secondary goal of this study was to compare expression of several neuroimmune molecules between males and females in the pursuit of potential molecular determinants of the observed sex differences in drinking behavior. We observed sex-specific time course responses of several immune genes, with females showing a higher increase and slower decline in expression of pro-inflammatory cytokines, which may, at least in part, explain sex differences in the immune modulation of alcohol consumption.

High alcohol consumption is a hallmark of AUD. The neuroimmune hypothesis of excessive EtOH drinking is based on the idea that consumption of large amounts of alcohol promotes pro-inflammatory signaling, which, in turn, results in more alcohol intake, creating a positive feedback loop that may lead to excessive alcohol consumption (Crews & Vetreno, 2016). Studies using chronic intermittent EtOH drinking paradigms show that mice escalate EtOH consumption and preference over time, which provides support for this hypothesis (Blednov et al., 2005, 2010; Hwa et al., 2011; McCarthy et al., 2018; Osterndorff-Kahanek et al., 2013; Ozburn et al., 2010; Ponomarev et al., 2017). In this study, we used FVB/B6 and B6/FVB F1 hybrid mice, which were previously shown to consume more EtOH than their respective parental strains (Blednov et al., 2005, 2010; Ozburn et al., 2010). Our results indicate that saline-treated female, but not male mice demonstrate a stable increase in EtOH drinking and preference over sessions. On the other hand, innate immune activation by PIC resulted in escalation of both consumption and preference in males, but not females. These sex differences may, at least, in part, be explained by a potential “ceiling” effect. Female mice drink considerably more EtOH per body weight and further escalate their drinking over time, potentially leaving no “room” for a further increase in consumption. These findings are consistent with other reports that used the B6 mouse strain (Warden, DaCosta, et al., 2020; Warden et al., 2019a, 2019b; Warden, Triplett, et al., 2020). Interestingly, compared to B6 findings, our data in F1 males show that TLR3-dependent immune activation results in a potentially higher escalation in EtOH drinking that occurs earlier during the 5-week drinking period.

We also tested if the mode of innate immune activation plays a role in modulation of EtOH drinking by using PIC, the TLR3 agonist and LPS, the TLR4 agonist. Compared to PIC, LPS resulted in a decrease in EtOH consumption and preference in FVB/B6 males and no change in females. These results contradict previous findings showing that LPS produced persistent increases in EtOH consumption in B6 and FVB/B6 mice (Blednov et al., 2011). There are many differences in experimental design between the two studies, which may potentially explain the contrasting findings. Compared to the Blednov et al study, we used a smaller dose of LPS injected repeatedly, which may have produced a different pro-inflammatory state. We also used a different concentration of EtOH, 15% v/v, compared to a range of different doses in the other study. The lack of the effects of LPS and PIC in females was consistent with previous reports in B6 mice using a similar paradigm (Warden et al., 2019a, 2019b).

Further evidence for the effects of genotype on the immune modulation of EtOH consumption could be obtained from recent studies. Warden and colleagues tested two similar inbred mouse substrains: C57BL/6J (B6J) and C57BL/6N (B6N) and showed that even small genotypic differences can result in robust behavioral changes, as only B6J males showed PIC-induced increase in EtOH intake. To further explore the effects of sex and genotype on the immune modulation of alcohol intake, we carried out a pilot experiment using Camk2axSun1 F1 hybrid male and female mice. The F1 mice are considered to be mainly on the B6J genetic background, but also include genetic material from other inbred strains including B6N. We used a similar behavioral paradigm and found that repeated injections of PIC did not change EtOH consumption in females, but had a strong trend to reduce alcohol drinking in male mice (Hitzemann et al., 2021). Other studies provided additional evidence for the role of genotype and species in the immune modulation of EtOH drinking (Blednov et al., 2011; Gano et al., 2022; Giffin et al., 2022; Lovelock et al., 2022; Warden, DaCosta, et al., 2020; Warden et al., 2019a, 2019b; Warden, Triplett, et al., 2020).

Chronic EtOH consumption results in changes in immune-related gene expression within the brain. Increases in pro-inflammatory gene expression have consistently been detected in post-mortem human brains from patients with AUD (Brenner et al., 2020; He & Crews, 2008; Lewohl et al., 2000; Liu et al., 2006; Ponomarev et al., 2012) and preclinical rodent models (Crews & Vetreno, 2016; Mulligan et al., 2006, 2011; McCarthy et al., 2018). It was previously suggested that sex and genotype differences in the immune modulation of EtOH consumption may, at least, in part, be explained by differences in temporal profiles of pro-inflammatory molecules in brain. Warden and colleagues measured time-dependent expression of pro-inflammatory genes and showed that the peak immune activation in B6J males occurs earlier than in B6J females (Warden et al., 2019a, 2019b). Based on their behavioral data, authors hypothesized that availability of EtOH during peak activation would result in decreased drinking or no changes, whereas alcohol consumption during the descending limb of immune activation may increase EtOH intake. Our molecular data provide partial support for this hypothesis. In our current study, we screened several known proinflammatory genes (*Il1b, Tnf, Ccl5*) which have been previously implicated in regulation of EtOH consumption (Bajo et al., 2019; Cooper et al., 2020; Crews et al., 2006; Gano et al., 2019; McCarthy et al., 2018; Nwachukwu et al., 2023; Roberto et al., 2018). We identified that mRNA expression of these pro-inflammatory molecules was significantly higher at the peak and lasted longer in females compared to males after the final 10th injection of PIC. These results are complimentary to other studies suggesting that immune activation by PIC is stronger in females compared to males (Chavez-Valdez et al., 2019; Hitzemann et al., 2021).

One of the main objectives of our study was to extend previous research on the immune modulation of EtOH consumption in mice to a genotype that has not been tested for the effects of TLR3 activation. We used the high EtOH-consuming FVB/B6 and B6/FVB F1 hybrid mice and showed that TLR3-dependent innate immune activation resulted in an escalation of EtOH drinking in males but not females. These effects in males were reproducible and consistent across different doses of PIC. Sex differences in the immune modulation of EtOH consumption may, at least, in part, be explained by the potential “ceiling” effect in drinking patterns and differences in temporal profiles of pro-inflammatory molecules in brain. Taken together, these results support previous findings that the effects of immune activation on EtOH consumption depend on genotype, sex, and mode of activation and suggest that FVB/B6J and B6J/FVB F1 males are a suitable model to study TLR3-dependent escalation of alcohol drinking.

## Acknowledgements

We would like to thank Luke Wagner for his technical assistance for the behavioral experiments. This research was funded by the National Institutes of Health/National Institutes on Alcohol and Alcoholism AA027096 and AA028370 to IP and American Heart Association 24PRE1184797 to BRK. Diagram schematics were generated using BioRender.

## Author Contributions

Brent R. Kisby: methodology, formal analysis, investigation, data curation, writing original draft, funding acquisition, review and editing.

Isabel Castro-Piedras: methodology, formal analysis, investigation, data curation, writing original draft, review and editing.

Sambantham Shanmuganum: methodology, writing original draft, review and editing.

Igor Ponomarev: methodology, formal analysis, investigation, data curation, review and editing, project administration, funding acquisition.

## Data availability

Request for data upon reasonable request

